# Deep Learning Approach to Genomic Breakage Study from Primary Sequence

**DOI:** 10.1101/2021.06.03.446904

**Authors:** Pora Kim, Hua Tan, Jiajia Liu, Mengyuan Yang, Xiaobo Zhou

## Abstract

Identifying the molecular mechanisms related to genomic breakage is an important goal of cancer mechanism studies. Among the diverse location of the breakpoints of structural variants, the fusion genes, which have the breakpoints in the gene bodies and typically identified from RNA-seq data, can provide a highlighted structural variant resource for studying the genomic breakages with expression and potential pathogenic impacts. In this study, we developed FusionAI which utilizes deep learning to predict gene fusion breakpoints based on primary sequences and let us identify fusion breakage code and genomic context. FusionAI leverages the known fusion breakpoints to provide a prediction model of the fusion genes from the primary genomic sequences via deep learning, thereby helping researchers a more accurate selection of fusion genes and better understand genomic breakage.

**Highlights:**

- FusionAI, a 9-layer deep neural network, predicts fusion gene breakpoints from a DNA sequence
- FusonAI reduce the cost and effort for validating fusion genes by decreasing specificity
- High feature importance scored regions were apart 100nt on average from the exon junction breakpoints
- High feature importance scored regions overlapped with 44 different human genomic features
- Transcription factor fusion genes are targeted by the GC-rich motif TFs
- FusionAI gives less scores to the non-disease derived breakpoints

## INTRODUCTION

Identifying the molecular mechanisms related to the genomic breakage is an important goal of disease biology to understand the origin of new genes and aberrant functional features of the broken genomes. Among the diverse location of the breakpoints of structural variants, the breakpoints of fusion genes are located in the gene bodies. Fusion genes can be formed mainly by double-strand breakages. Due to the cost-effective sequencing (data creation) and analyzing (interpretation), and usage (diagnosis), there are a huge amount of RNA-seq data accumulated to date. Fusion genes identified from RNA-seq data, the expressed ones, can provide a highlighted structural variant resource for studying the genomic breakages with expression and thereby potential pathogenic impacts. Indeed, the broken gene context of the identified fusion genes provided the aberrant functional clues to study disease genesis. However, an inherent limitation of RNA-seq data and analyses is restricted by sequencing depth, read length, read alignment tool and their options, filtering criteria, and etc., which create many false positives and cannot identify fusion genes with low expression. Therefore, developing the sequencing-free prediction of fusion genes would be helpful and may provide new insights into the genomic breakage phenomenon.

Motivated by recent successes in the use of deep learning approaches to predict the genomic regulatory elements and splice events from the genomic context (Jaganathan et al., 2019; Zhou and Troyanskaya, 2015), we hypothesized that the exon junctional breakpoints of known fusion genes from the RNA-seq data can be used to construct a deep-learning model of predicting the breakage tendency. To test this hypothesis, we developed FusionAI, a deep residual neural network that predicts whether an exon junction position pair of a potential fusion gene is a fusion gene breakpoint or not, using as input only the primary sequence. FusionAI consists of a deep neural network (DNN) model that classifies between fusion-positive and - negative breakpoints on the basis of DNA primary sequences. We used the fusion genes that have exon junction-junction breakpoints in both fusion partner genes of The Cancer Genome Atlas (TCGA)(Cancer Genome Atlas Research et al., 2013) cohorts annotated from FusionGDB(Kim and Zhou, 2019). We also made fusion negative breakpoint sequences by stringent criteria and we trained and tested the FusionAI model by mixing these fusion-positive and -negative data. We evaluated the performance of FusionAI by applying multiple identified fusion gene resources such as validated 2200 fusion genes from 675 human cancer cell-lines(Klijn et al., 2015), Sanger sequencing-based fusion genes in Entrez from ChiTaRs-3.1(Gorohovski et al., 2017), fusion genes of a non-disease population of Genotype-Tissue Expression (GTEx)(Consortium et al., 2017; Singh et al., 2020), and fusion genes derived from the structural variants of The Genome Aggregation Database (genomAD)(Collins et al., 2020) annotated from FGviewer(Kim et al., 2020). The application of FusionAI increased the specificity of fusion prediction. We found the relationship between feature importance (FI) score and FusionAI prediction scores for classifying fusion-positives and -negatives. From the top 10% (high) FI score regions across 20 kb long DNA input sequence, we investigated the human genomic sequence features. We identified significant overlap between 44 different human genomic sequence features and the regions near to the fusion gene breakpoints in diverse categories of features such as virus integration sites, multiple types of repeats, structural variant regions, specific chromatin stated regions, and expression regulating regions. In these top 10% FI scored regions of the 5’-gene breakpoint area of the transcription factor fusion genes, we identified a GC high DNA sequence motif, which might be targeted by SP1. The low percentage of FusionAI prediction scores in the healthy population derived fusion genes might reflect a validity or tumorigenicity of individual fusion gene breakpoint. In summary, FusionAI is an example of interpretable scientific deep learning in studying the human genomic breakages with diverse potential genomic regions related to different cellular mechanisms.

## RESULTS

### Overview of FusionAI

First, we investigated the distribution of the breakpoint location of fusion genes on the exonintron gene structure. The majority of the fusion genes derived from RNA-seq data were located at the exon junction-junction breakpoints (Figure 1A). This confirms the hypothesis that the breakpoints of fusion genes would be located in the intronic region since the exons cover the human genome only about 1%, but the introns cover more than 24% except the intergenic regions (Venter et al., 2001). This percentage will be bigger in the view of gene body. From this context, we mainly used the fusion gene breakpoint information located at the exon junction-junction among TCGA fusion genes as the fusion-positive data sets (~26k fusion breakpoints in Table S1). We made a similar number of fusion-negative data with strict criteria (Figure 1B. See the methods section). Using the divided data from the mixture of fusion-positives and -negatives into 80% and 20%, we trained and test FusionAI. The input is a total 20 kb DNA sequence of combined ones of both fusion partner genes with 5kb +/- breakpoints. The transformed one-hot encoded input resultant into the probability of fusion breakpoints through passing the deep learning processes including filtering, activation, pooling, flattening, and fully connected functions (Figure 1C). To examine long-range and short-range specificity determinants, we compared the scores assigned to fusion-positives by the models trained on 200nt, 1k, 2k of the sequence context versus the full model that is trained on 5k of context (Figure 1D). Overall, there was no big difference in the accuracy across the different sequence range from the fusion breakpoints. The accuracies for training and test data sets were 97.4% (AUCROC=0.9962) and 90.8% (AUCROC=0.9706) with 0.12 and 0.42 error rate, respectively. This performance is much better than the traditional machine learning method s, SVM and RF. To identify the hidden genomic context features specific fusion gene breakpoints, we performed our studies based on the 5k based model with a convolutional neural network approach.

**Figure 1.**
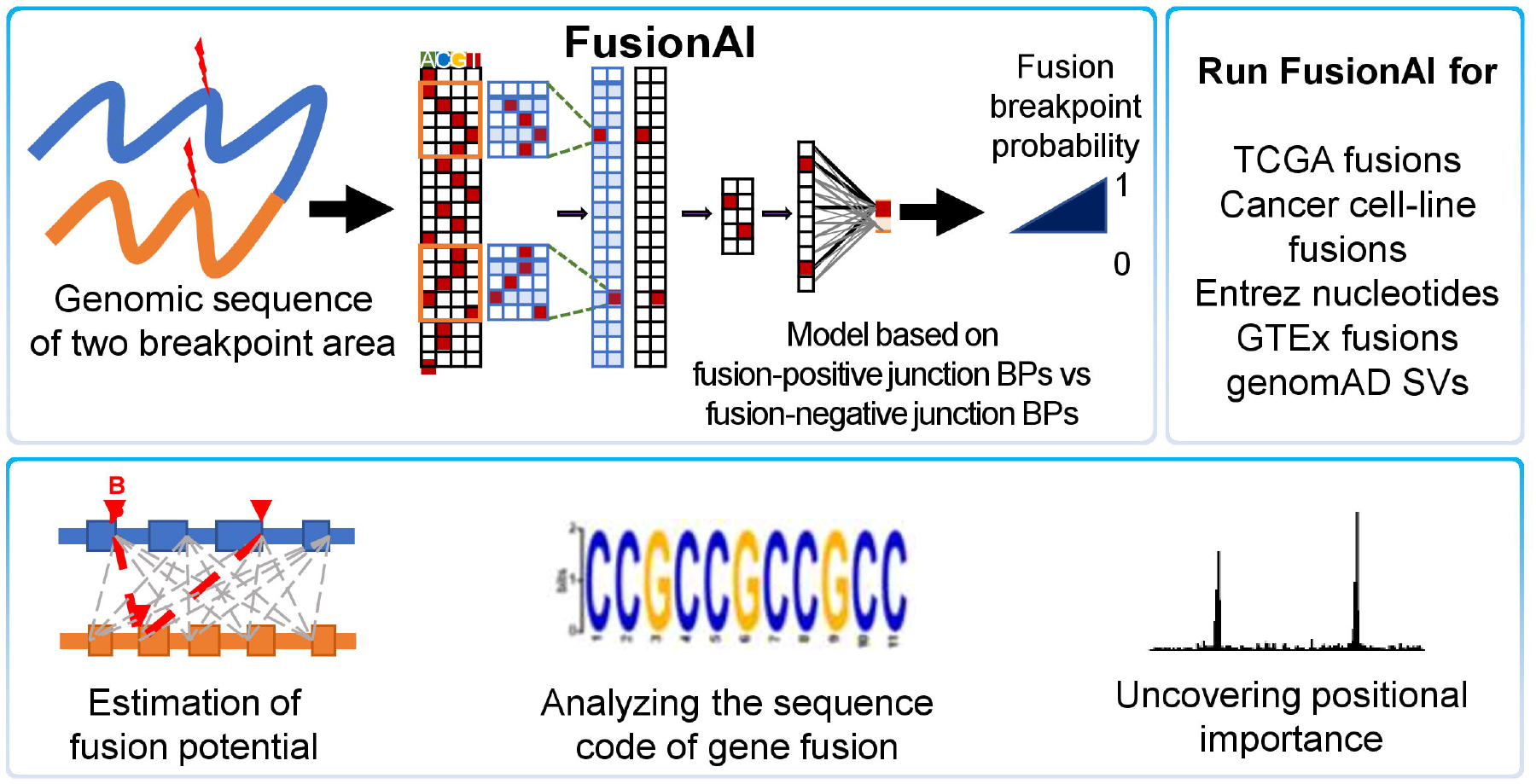
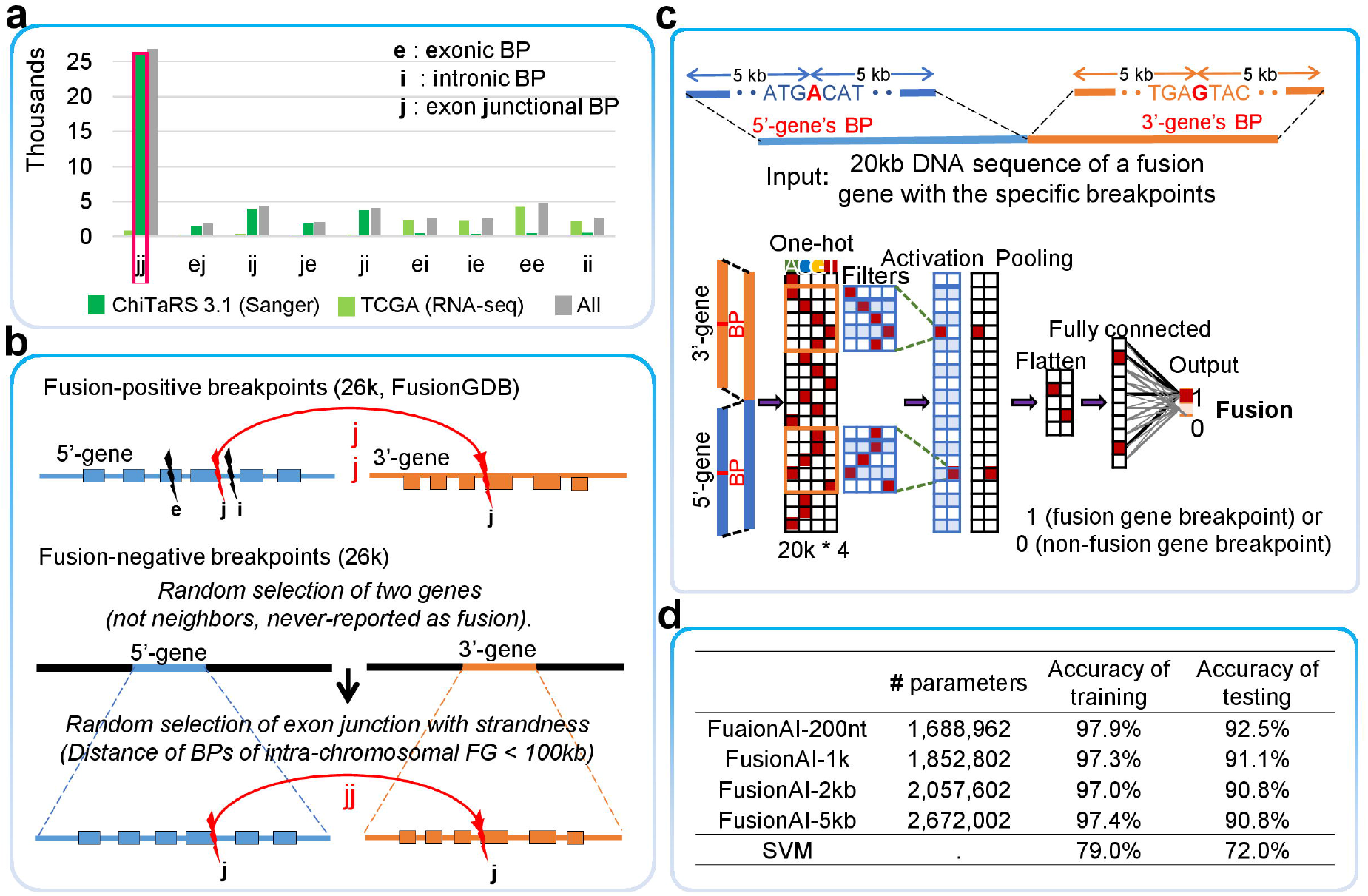
Overview of FusionAI. (a) The investigation of fusion gene breakpoints of 48K FGs from FusionGDB identified the BP location across the human genome. (b) Making training and test dataset of fusion-positive and -negative breakpoints. (c) Diagram of fusion gene breakpoints classification by FusionAI. (d) Effect of the size of the input sequence context on the accuracy.

### Primary sequence based FusionAI improves identification of fusion genes

To compare the performance of prediction of fusion genes of FusionAI, we chose STAR-fusion and Arriba based on the best performance results from the accuracy assessment study (Haas et al., 2019). Our training and test data sets are made of the genomic breakpoint information. Since the usual RNA-seq based tools require the input of RNA-seq data, we made the simulation RNA-seq data of the split reads at the exon junction breakpoints with different read length (50, 75, and 100 bp) and a different number of split reads (1, 3, 5 split reads, and 10 random around breakpoints) based on the fusion-positive and -negative breakpoints in our training and test data sets (Figure 2A). We also made simulation RNA-seq data for the experimentally validated fusion gene breakpoints by Sanger sequencing and RT-PCR from ChiTaRS-3.1(Gorohovski et al., 2017) and cancer cell-line study(Klijn et al., 2015), respectively. To apply FusionAI to these data sets, we made the input sequence of 20kb long (Table S1). As shown in these comparisons in FigureS 2A and 2B, the typical RNA-seq’s limiting factors (i.e., read-length and the number of split reads) were not problems to FusionAI compared to the general RNA-seq based fusion prediction tools as shown in the red triangles in the plots (Figures 2A and 2B, Tables S2-4). We also checked the individual fusion genes validated from the most famous fusion gene-positive cell-lines such as K562, MCF7, and NCI-H660, which are the *BCR-ABL1, BCAS4-BCAS3*, and *EML4-ALK-positive* cells, respectively (Figure 2C). Figure 2D shows that FusionAI has the biggest number of validated fusion genes among the three tools. Typically, the researchers use multiple fusion gene prediction tools to select the candidates before experimental validation. Compared to the result using only two tools of STAR-fusion and Arriba, additional use of FusionAI reduced the number of false positives effectively, but remain the true positives. This can reduce the cost and efforts for validation.

**Figure 2.**
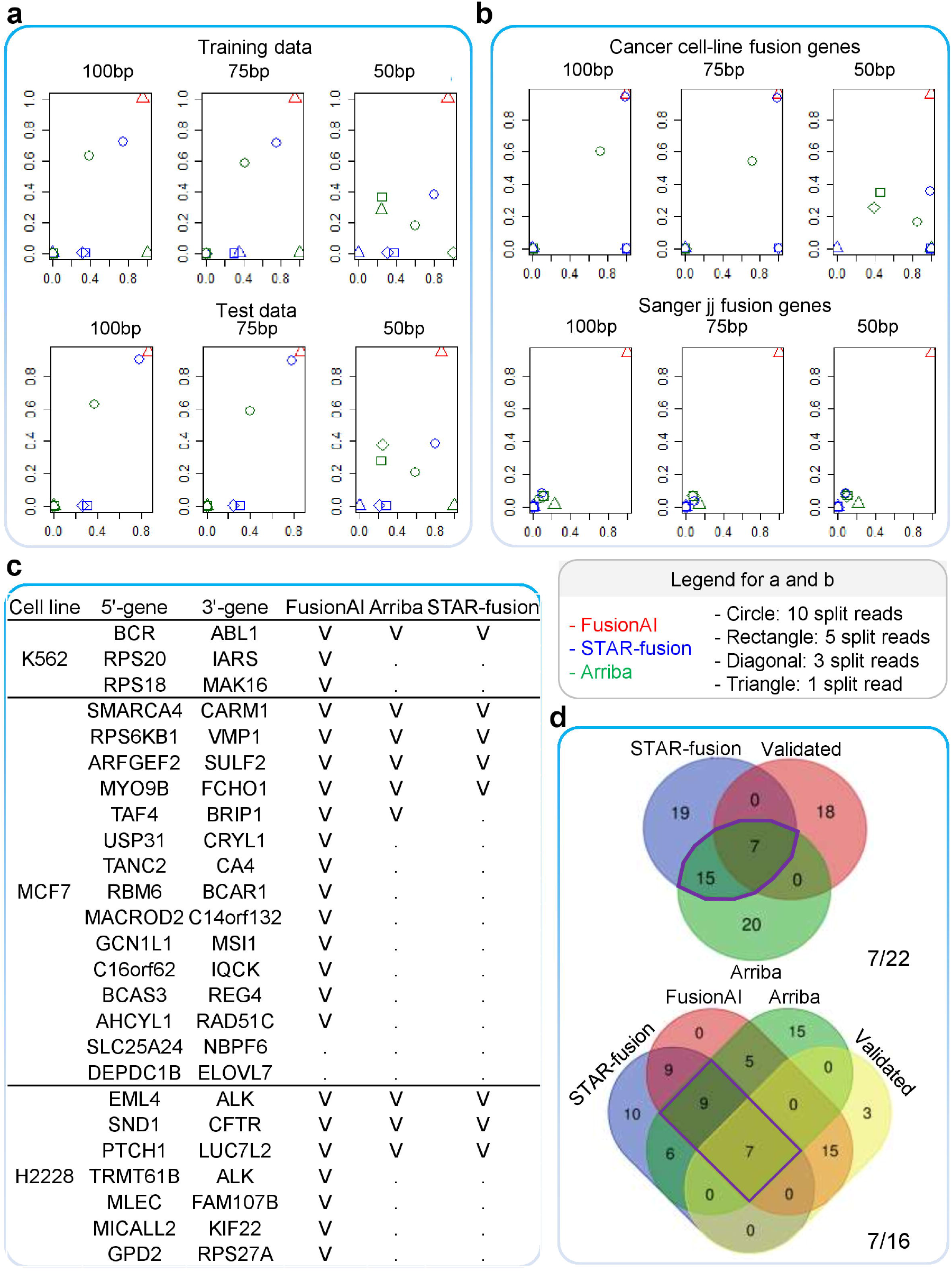
Performance of FusionAI. (a) Comparison of FusionAI to other methods for fusion gene prediction including 38,000 TCGA fusion genes from training and test data sets. The plots show the number of true positives and sensitivity from left. (b) Comparison of the number of the true positive fusions in ~ 2200 validated fusion genes in ~530 cancer cell-lines, and 862 Sanger sequence based fusion genes that have fusion breakpoints at the exon junction position from ChiRTaRS3.1. (c) Comparison of predicting fusion genes in three cancer cell-lines (H2228, K562, and MCF7). (d) Identification of validated fusion genes in three cell-lines.

### FusionAI facilitates better understanding of genomic context of fusion gene breakpoints

We investigated the FI scores using 20bp window size every 20bp along the 20k bp input sequence of our training and test fusion genes. These FI scores are the values showing how big impact of individual 20bp length sequences along 20k have to distinguish fusion-positive and - negative breakpoints. As shown in the six most famous fusion genes (*BCR-ABL1, EML4-ALK, TMPRSS2-ERG, PML-RARA, RUNX1-RUNX1T1, and FGFR3-TACC3*), the FI scores were most high at the breakpoint area overall (Figure 3). For the intuitive vaalidation, we checked the performance of FusionAI how much the output scores classify the fusion-positives and - negatives using the logistic regression method. As shown in Figure 3B, FusionAI classifies the fusion-positive and -negative breakpoints very significantly (p-value <2e-18). We also wondered about relationship between the FI scores and the FusionAI output scores. Because some fusion genes had very small values of FI scores (i.e., *BCR-ABL1* in Figure 3A). We transformed the original values of FusionAI and FI scores to the quantile normalized values. Then, we identified existence of grouping tendency between fusion-positives and -negatives among the FusionAI output scores, and second and third principal components from the principal component analysis as shown in Figure 3C.

**Figure 3.**
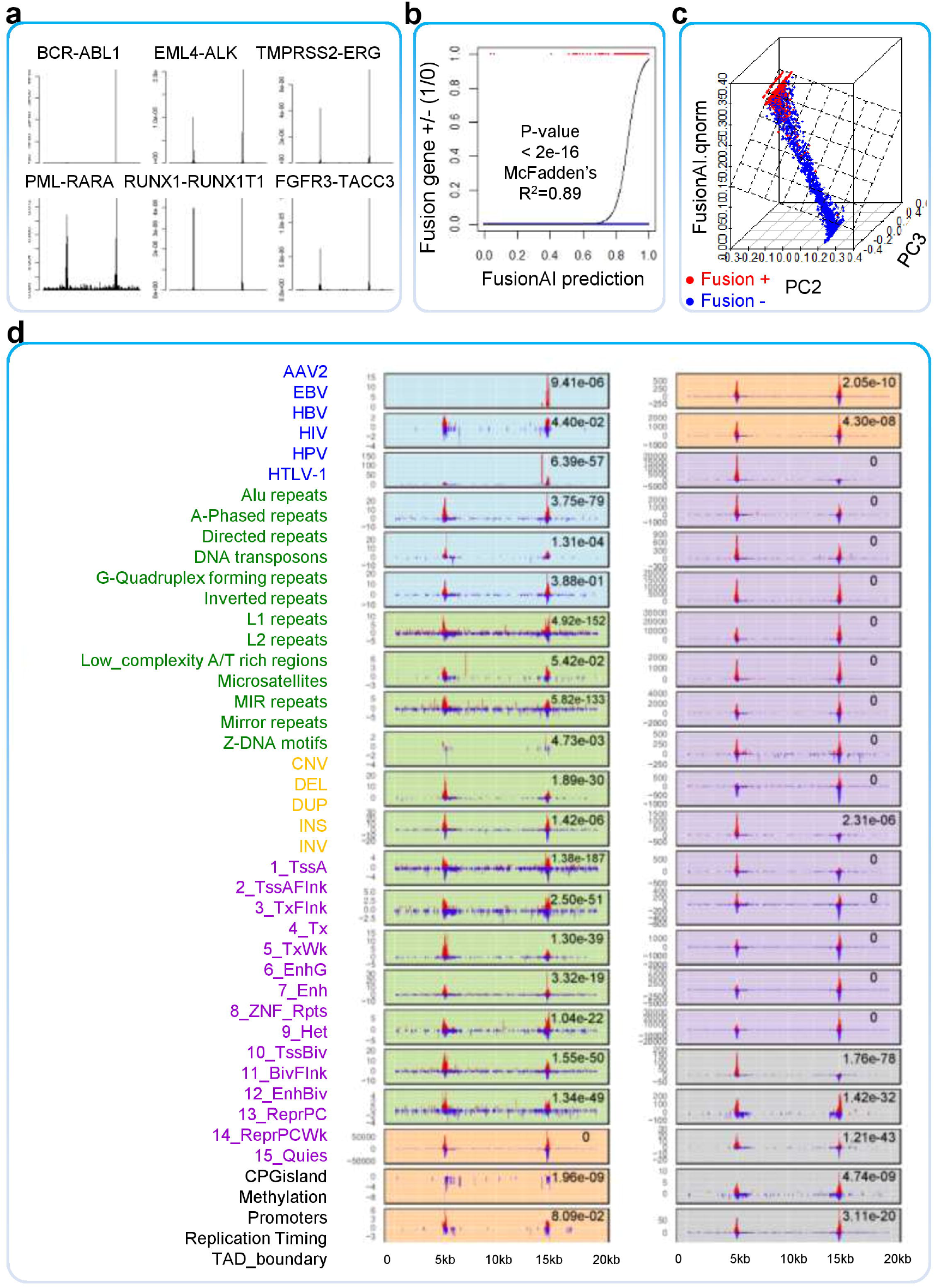
Feature importance score for understanding genomic breakage. (a) Distribution of FI scores across 20kb long of six representative fusion gene breakpoints. (b) Logistic regression result of FusionAI prediction. (c) Classification between fusion-positive and -negative from FusionAI and FI scores. (d) Distribution of overlapping between top 10% FI scored regions and 44 different types of human genomic features in 5 different categories.

### High feature importance scored regions provide a landscape of the genomic feature aspect of fusion gene breakpoints

To date, there were many trials to understand the gnomic features of breakage to study multiple effects of the genomic breakage(Ballinger et al., 2019; Chakraborty et al., 2020; Fungtammasan et al., 2012; Peng et al., 2006). In this study, we sought to identify fusion-positive breakpoint-specific genomic features across human genome sequence by integrating 44 different human genomic features of five categories such as integration sites of 6 viruses, 13 types of repeats, 5 types of structural variants, 15 different chromatin stated regions, and 5 gene expression regulatory regions. For individual feature categories, we counted the unique number of the overlapped loci of the feature with the top 10% FI scored regions in every nucleotide across 20k sequence in both fusion-positive and -negative groups. As shown in the first six plots of Figure 3D, the overall distribution of overlapping was enriched in the fusion gene breakpoint area. Moreover, these distributions were significantly different between fusion-positive and -negative breakpoint groups (Table S5). The overall distance between the high FI scores and breakpoints was 70 nucleotides median, 99.54 nucleotides mean with 211.28 standard deviations as shown in Figure 4A. This would explain the outperform of the FusionAI-200nt model among others. More interpretations on the individual features are described below.

**Figure 4.**
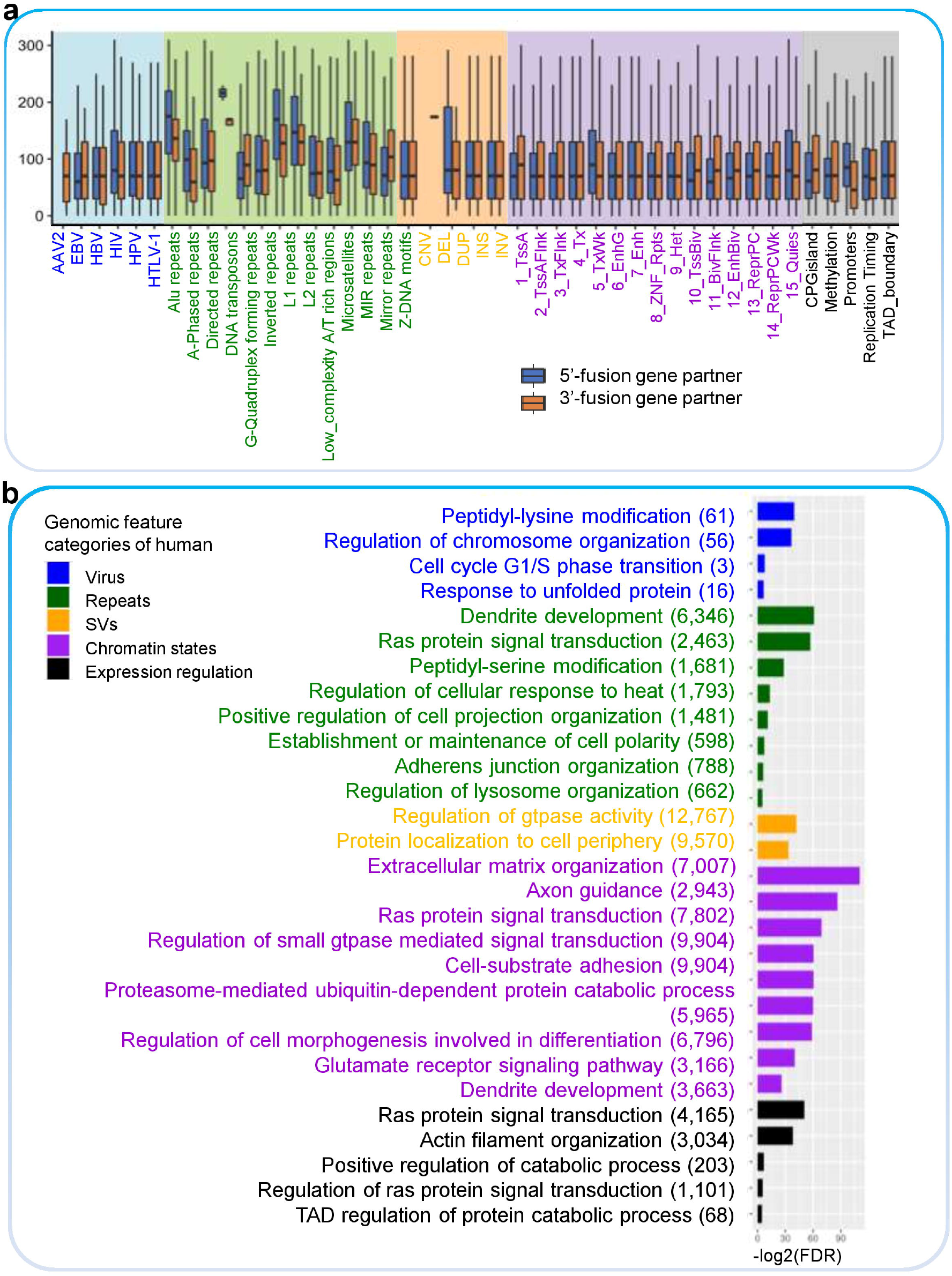
High feature importance scored regions. (a) Distribution of the distance between the high FI scored regions and the exon junctional breakpoints. (b) Enriched biological processes in the genes that have overlap with high FI scored regions per individual genomic feature categories.

#### Virus integration sites

Specifically, fusion-positive breakpoints were enriched in the virus integration sites of hepatitis B virus (HBV) and human immunodeficiency virus (HIV). Gene ontology enrichment test identified that the high FI scored region genes overlapped with the HIV integration sites were enriched in ‘peptidyl-lysine modification’ and ‘regulation of chromosome organization’. (Figure 4B). Multiple post-translational modifications (PTMs) of viral and cellular proteins gain increasing attention as modifying enzymes regulate virtually every step of the viral replication cycle (Chen et al., 2018). For HBV, ‘cell cycle G1/S phase transition’ and ‘response to unfolded protein’ were the enriched biological pathways. A hallmark of chronic hepatitis B virus (HBV) infection is known as containing excessive hepatitis surface antigen (HBsAg) in the endoplasmic reticulum (ER), which is linked to unfolded protein response (UPR) (Li et al., 2019). A previous study showed that HBV-infected primary human hepatocytes are enriched in the G2/M phase compared to the predominantly G0/G1 phase of cultured primary human hepatocytes (Xia et al., 2018).

#### Repeats

Next, fusion-positive breakpoints were enriched in multiple types of repeats as shown in green background plots of Figure 3D. Among 13 types of repeats, Alu repeats, direct repeats, and L1 repeats showed very significantly different distribution between fusion-positive and - negative groups. Alu elements are the most abundant transposable elements, containing over 1.2 million copies, which comprise 11% of the human genome (Deininger, 2011). It is reported as enriched in the common fragile sites (Fungtammasan et al., 2012). Direct repeats are known as eliciting genetic instability by both exploiting and eluding DNA double-strand break repair systems in mycobacteria (Wojcik et al., 2012). The human LINE-1 retrotransposon is known to create DNA double-strand breaks (Gasior et al., 2006). There was no specific difference in the structural variants compared to other features. This might be explained by that our training data set is made up with the RNA-seq based exon junction breakpoints, which is different from whole-genome sequencing-based structural variants.

#### Chromatin states

The purple background plots are the overlapping result with the chromatin state calls, using a 15-state model from Roadmap Epigenomics Mapping Consortium (Roadmap Epigenomics et al., 2015). As shown in the first four plots of the purple background ones in the right panel of Figure 3D, the top 10% FI scored fusion-positive breakpoint area of 5’-genes enriched in the active chromatin states, which were associated with the expressed genes such as active transcription start site (Tss) proximal promoter states (TssA, TssAFlnk), a transcribed state at the 5′ and 3′ end of genes showing both promoter and enhancer signatures (TxFlnk), actively transcribed states (Tx). However, the fusion-negatives were relatively more enriched with the high FI scored regions related to the repressed chromatin states. In other words, the breakpoints of fusion-positive breakpoints are located at the transcriptionally active chromatin states’ peak regions, but the ones of fusion-negatives are located at the transcriptionally repressed chromatin states’ peak regions. This pattern might be related to the typical roles of the driver fusion genes as the transcriptional activation itself or downstream target genes (Kim et al., 2017; Kim et al., 2018; Kim et al., 2020). This makes sense since the 5’-gene partner’s promoters will be used as the promoter of the fusion genes.

#### Gene expression regulatory

The first plot in the last category of gene expression regulatory with gray color background shows the more breakpoints of 5’-genes (3,684) are located in the CpG island area than 3’-genes (678). This might provide additional evidence for the previous finding that initial chromosomal breakage occurs directly at or near CpGs (Tsai et al., 2008). 4,165 genes in these regions were enriched in the ‘Ras protein signal transduction’ pathway. Imbalance of the Ras signaling pathway is a major hallmark of human cancer (Irimia et al., 2004). Furthermore, 3,849 out of 4,165 genes were mainly targeted by MAX interactor 1, dimerization protein (MXI1), and enhancer of zeste 2 polycomb repressive complex 2 subunit (EZH2). MXI1 is the MYC antagonist, also regarded as a tumor suppressor. EZH2 is the enzymatic subunit of Polycomb repressive complex 2 (PRC2), a complex that methylates lysine 27 of histone H3 (H3K27) to promote transcriptional silencing (Kim and Roberts, 2016). EZH2 is known to regulate MXI1 by targeting from the harmonizome, a collection of processed datasets gathered to serve and mine knowledge about genes and proteins (Rouillard et al., 2016). Through the DNA breakage of fusion genes, these CpGs located genes might be expected to have aberrant regulation governed by EXH2. The last feature, replication timing has been notified as being associated with the nature of chromosomal rearrangements in cancer (Du et al., 2019). Similar to CpGisland, there was 1,101 genes enriched in the ‘regulation of ras protein signal transduction’ pathway. In the methylation and TAD_boundary features, 203 and 68 genes were enriched in the ‘positive regulation of catabolic process’ and ‘regulation of protein catabolic process’. Overall, this analysis identified enriched biological pathways of ras signal transduction and cellular regulation of catabolic processes in the gene expression regulatory that share the genomic breakages. From this, we can infer the involvement of genomic breakage in the cancer cell proliferation or maturation, and metabolic reprogramming.

#### The GC-rich motif is enriched in the 5’-genes’ breakpoint area of transcription factor fusion genes

We wondered what DNA motif sequences would work in high FI scored regions, which are located about 100nt +/-200 SD apart from the exon junction breakpoint. We checked the high FI score regions of all fusion-positive breakpoints of training data. Then, we identified the top two most frequent motifs – TTMWTTTTTTTTTTTTTTYYT and CGGCGGCGGCGGCGGCGGCGG using MEME Suite (Bailey et al., 2015) (Figure 5A). The genes that have the former motif in their promoter regions were enriched in these biological processes, ‘Regulation of organ growth’, ‘Negative regulation of alpha-beta T cell differentiation’, and ‘mRNA metabolic process’. Because 2 out of 3 known fusion genes are the intra-chromosomal rearrangement events, we searched DNA motif sequences focusing on these events. Then, there were the top two T-rich motifs, which were enriched in these biological processes - ‘transcription initiation from RNA polymerase II promoter’, ‘G-protein coupled receptor protein signaling pathway’, ‘negative regulation of leukocyte activation’, ‘response to stimulus’. To date, there are two types of the most popular driver fusion genes like kinase fusion genes and transcription factor fusion genes as helping cancer cells’ proliferation and progression (Kim et al., 2017; Kim et al., 2018). We identified ‘HTTTTBTTTTT’ motif around high FI score regions in the fusion genes composed with dimerization and kinase domain in the individual partner genes. The genes with this motif in the promoter region were involved in the ‘cellular macromolecule biosynthetic process’. From the transcription factor fusion genes, we found a relatively long (28 nt) GC-rich motif, ‘GCBGGSSGSGGSSGSSSGGGGSGSBGGG (Supplementary Table S2). The genes with this motif in their promoter region mainly involved in ‘Negative regulation of signal transduction’. For example, Figure 5B shows the distribution of this GC-rich motif sequence across 20kb sequence of *TMPRSS2-ERG* fusion gene. Interestingly, this GC-rich motif sequence was located around the breakpoint of the 5’-gene. *TMPRSS2-ERG* is known as dependent on the expression of Sp1 (Meisel Sharon et al., 2016). Sp1, a transcription factor, binds to GC-rich motifs (Koutsodontis et al., 2002). Figure 5C shows the recurrent (expressed in more than 3 samples) transcription factor fusion genes. Specifically, 17 fusion genes out of these 22 were bound by Sp1 as shown in the searching result from ENCODE transcription factor target database and TRRUST (Consortium, 2011; Han et al., 2018). Total 220 genes in the transcription factor fusion genes have this GC-rich motif. Out of these, 51 genes were transcription factors and 38 genes were epigenetic factors. Unique 80 genes of 51 plus 38 were annotated as mainly involved in the diverse epigenetic mechanisms as shown in Figure 5D such as ‘DNA methylation and alkylation’, ‘histone modification’, ‘transcription regulation’, ‘cellular response to ROS’, and etc. In this study, we may have evidence of that the transcription factor fusion genes might regulate their downstream targets by using the epigenetic mechanisms.

**Figure 5.**
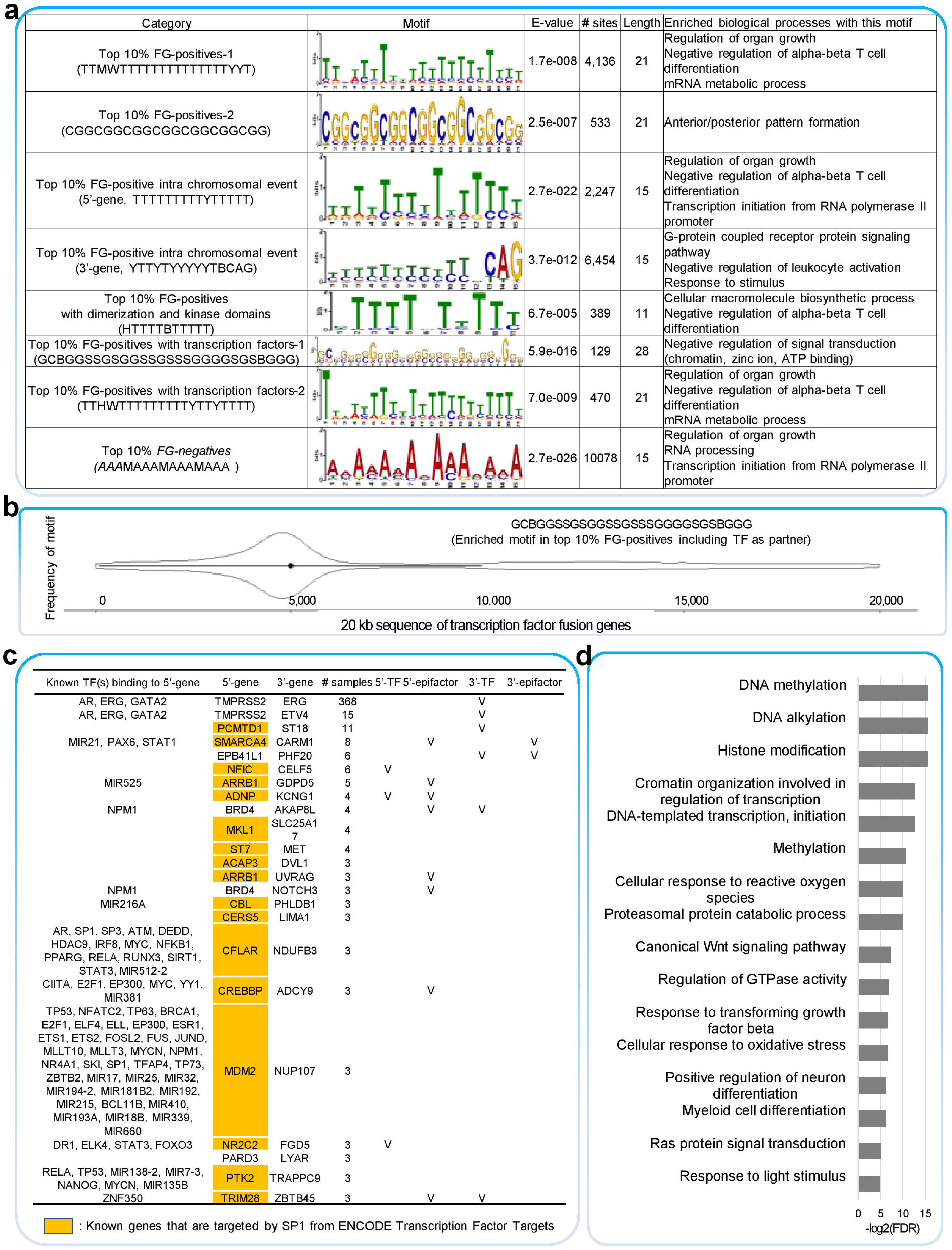
Consensus motif sequences in the high FI scored FG-positive regions and enriched biological processes. (a) Identified DNA sequence motifs in fusion-positive breakpoint area of multiple groups such as all fusion-positives, intra-chromosomal events of fusion-positives, kinase fusion genes with dimerization and kinase domain, transcription factor fusion genes. (b) Distribution of the GC-rich motif across 20kb length sequence in the isoforms of TMPRSS2-ERG fusion gene. (c) Transcription factor fusion genes that have GC-rich motifs. (d) Enriched biological processes of those genes that have GC-rich motifs.

### Chimeric transcripts from the non-disease tissues have low FusionAI score

Fusion genes formed by the chromosomal rearrangements are the hallmark of cancers from genomic instability. However, chimeric transcripts are also existing in non-cancerous cells and tissues (Babiceanu et al., 2016; Finta and Zaphiropoulos, 2002; Li et al., 2009; Yuan et al., 2013). We wondered how FusionAI output scores would reflect this difference of the sample sources of fusion genes. To compare the prediction of FusionAI from diverse cohorts, we integrated the fusion genes whose breakpoints are located at the exon junction-junction loci from TCGA, cancer cell-lines, and Sanger transcripts from Entrez, GTEx, genomAD (Table 1 and Table S2). We compared the ratio of the fusion genes whose FusionAI score is greater than or equal to 0.5. Here GTEx representing the healthy population was used to see how FusionAI prediction is different between cancerous and non-cancerous samples. Overall, fusion genes of the cancer cohorts, the top four groups in Table 1, had FusionAI output scores bigger than 0.5 in 93% of fusion gene breakpoints. However, if the fusion gene was common or exists in a healthy population only, then the ratio of the fusion genes with exon junctional breakpoints as predicted fusion-positive by FusionAI was decreased to 83% and 64%, respectively. Currently, we do not fully understand the meaning of this different ratio, but we guess this difference might be reflecting a validity or tumorigenicity of individual fusion gene breakpoint. genomAD, the whole genome-based structural variant data from diverse clinical cohorts, was used to see how FusionAI prediction is different between RNA-seq or WGS based fusion genes. From the previous study for visualizing the functional features of fusion genes at four different levels, FGviewer, we found 1,037 potential fusion genes whose breakpoints of the structural variants detected from genomAD v2.1 of 15K population WGS data were located on the gene bodies. Out of 1,037 fusion genes, 823 were the cases that have the exon junction-junction breakpoints. Among these, only 46 cases have been predicted as fusion-positive candidates by FusionAI (5%). Here, we used fusion genes anticipated as having exon junction-junction breakpoints from structural variants for genomAD. The small ratio might be from the different sequencing types of data and not expressed as the fusion transcript.

**Table 1.**
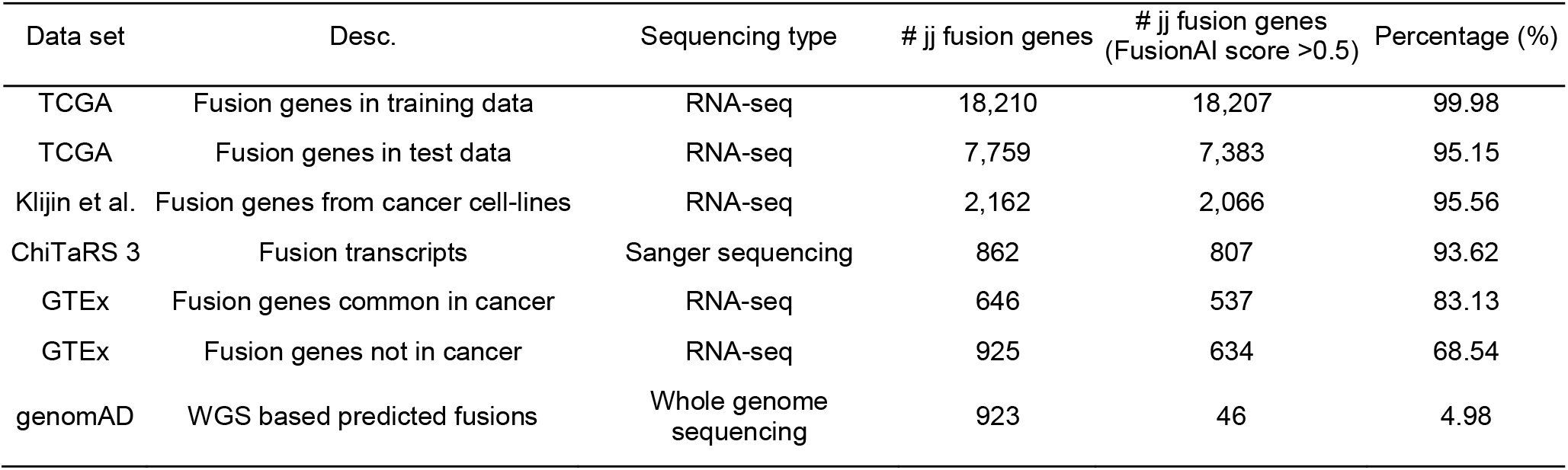
Comparison of FusionAI scores among the fusion genes in pan-cancer and healthy tissues.

| Data set | Desc. | Sequencing type | # jj fusion genes | # jj fusion genes<br>(FusionAI score >0.5) | Percentage (%) |
| --- | --- | --- | --- | --- | --- |
| TCGA | Fusion genes in training data | RNA-seq | 18,210 | 18,207 | 99.98 |
| TCGA | Fusion genes in test data | RNA-seq | 7,759 | 7,383 | 95.15 |
| Klijn et al. | Fusion genes from cancer cell-lines | RNA-seq | 2,162 | 2,066 | 95.56 |
| ChiTaRS 3 | Fusion transcripts | Sanger sequencing | 862 | 807 | 93.62 |
| GTEX | Fusion genes common in cancer | RNA-seq | 646 | 537 | 83.13 |
| GTEX | Fusion genes not in cancer | RNA-seq | 925 | 634 | 68.54 |
| genomAD | WGS based predicted fusions | Whole genome<br>sequencing | 923 | 46 | 4.98 |

## DISCUSSIONS

Our study suggests that by combining deep-learning predictions with empirical evidence in user-specific RNA-seq data, FusionAI predicts fusion gene breakage with high specificity and expands the findings beyond a conventional RNA-seq fusion gene analysis. From the study of high FI scored regions, we found that the overall distance between the high FI scores and breakpoints was 70 nucleotides median, 99.54 nucleotides mean with 211.28 standard deviations. This might explain one of the reasons why the FusionAI-200nt model outperformed other models using different flanking sequence length.

Before finalizing FusionAI model using the exon junction-junction breakpoints of both fusion-positive and -negative groups, we tried to build the initial version model by comparing the exon junction-junction breakpoints of 20K fusion-positives versus any breakpoint positions in the gene body of 20K fusion-negatives with +/- 5kb flanking sequences of each fusion partner gene. This initial model showed better performance than the current finalized model like the accuracy of 99.7% and 98.2% for training and test data, respectively. However, we recognized a potential issue regarding on the design of the comparison since this model can highlight and give more weights on the exon junction regions (exon-intron boundary) compared to the intronic breakpoints. The learned features from initial version model might mostly be the ones that are significantly related to the exon junctions like splicing signals. To avoid this wrong conclusion, we redesigned our modeling using only exon junction-junction breakpoints for both fusion-positive and -negative data and finalized FusionAI. Even though we are comparing the same conditions of exon junction-junction data, FusionAI found fusion-positive breakpoints well. In the future, enough validated data of the chimeric transcripts with specific breakpoints formed by the trans-splicing mechanism in the RNA-level would better explain the detailed exon junctional features that we found in this study.

Our trial for comparing the fusion gene breakpoints of different disease status or sequencing data found that FusionAI prediction gives higher scores for the cancerous and RNA-seq bases exon junction breakpoints. From this context, if we would have more fusion gene list from the diverse disease cohorts not only cancers, we may expect FusionAI to identify certain disease specific genomic breakage features by comparisons. Moreover, if there are plentiful number of fusion gene breakpoints at the DNA level that will be accumulated in the future, we will be able to build the genomic level breakpoint-based model and can bring deeper insight into the breakage mechanisms. Having enough amount of the real genomic breakpoints of the massive fusion genes to perform artificial intelligence approaches will need long time and many efforts. We hope we can make more precise new models based on the genomic breakpoint-based training data not by exon junctional breakpoints in the future. We believe it would be possible soon since the cost of WGS is decreasing down as time goes by with the maturation of the development of overall next generation sequencing technology and analysis methods.

Our study used the deep learning method to better understand the genomic breakage context focused on the fusion genes, which are the highlighted ones as expressed structural variants. Much work remains to be done to understand the human genome breakage in diverse disease, greater understanding of the genomic breakage mechanisms could pave the way for novel candidates for therapeutic intervention. Our findings using FusionAI model would enhance our understanding of fusion gene context. We hope FusionAI could serve as the initial platform of the efficient investigation of genomic breakage events.

## STAR * METHODS

Detailed methods are provided in the online version of this paper and include the following:

- KEY RESOURCES TABLE
- CONTACT FOR REAGENT AND RESOURCE SHARING
- EXPERIMENTAL MODEL AND SUBJECT DETAILS
- METHOD DETAILS
- DATA AND SOFTWARE AVAILABILITY

## SUPPLEMENTAL INFORMATION

Supplemental Information includes two figures and seven tables and can be found with this article online at.

## ACKNOWLEDGEMENTS

This work was partially supported by the National Institutes of Health grants [NIH R01GM123037, U01AR069395, and R01CA241930] to X. Zhou and [R35GM138184] to P. Kim. The funders had no role in study design, data collection, and analysis, decision to publish, or preparation of the manuscript. Funding for open access charge: Startup Fund to Dr. Kim from the University of Texas Health Science Center at Houston.

## AUTHOR CONTRIBUTIONS

P.K. arranged the training and test data by making fusion-positive and -negative data. H.T. performed the deep learning analysis. J.L. integrated human genomic feature information. P.K. made simulation RNA-seq data. M.Y. ran other fusion prediction tools. P.K. performed downstream analyses of DNA sequence motif and annotated findings. P.K., H.T., and X.Z. supervised the analyses and designed the project. P.K. wrote the paper.

## KEY RESOURCES TABLE

| REAGENT or RESOURCE | SOURCE | IDENTIFIER |
| --- | --- | --- |
| Deposited Data |  |  |
| Open reading frame annotation of known fusion genes | Kim et al. 2019 | https://ccsm.uth.edu/FusionGDB/ |
| Fusion gene breakpoint information of TCGA fusion genes | Kim et al. 2019 | https://ccsm.uth.edu/FusionGDB/ |
| Fusion gene breakpoint information of cancer cell-lines | Klijn et al. 2015 | N/A |
| Fusion gene breakpoint information of Sanger sequencing | Gorohovski et al. 2017 | http://biodb.md.biu.ac.il/chitars.prv |
| Fusion gene breakpoint information of GTEx cohorts | Singh et al. 2020 | N/A |
| Fusion gene breakpoint information of genomAD cohorts | Kim et al. 2020 | https://ccsmweb.uth.edu/FGviewer |
| Simulation RNA-seq data of all validation sets | This study | https://ccsm.uth.edu/FusionGDB/FusionAI/ |
| Fastq files for RNA-seq of K562 |  | Sequence Read Archive accession: SRR521460 |
| Fastq files for RNA-seq of MCF7 |  | Sequence Read Archive accession: SRR064286 |
| Fastq files for RNA-seq of H2228 |  | Sequence Read Archive accession: DRR016705.1s |
| Virus integration site information | Tang et al. 2020 | https://bioinfo.uth.edu/VISDB |
| Repeatmasker | Bao et al. 2015 | http://www.repeatmasker.org |
| MicroSatellite DataBase (MSDB) | Avvaru et al. 2020 | https://data.ccmb.res.in/msdb |
| Structural variant breakpoint information of genomAD | Lappalainen et al. 2013 | https://www.ncbi.nlm.nih.gov/dbvar |
| Chromatin state calls using a 15-state model | Roadmap Epigenomics Consortium et al. 2015 | N/A |
| Location of CpGisland, Methylation, Promoters | Lizio et al., 2019 | https://fantom.gsc.riken.jp/5 |
| Replication timing-specific peak regions | Davis et al., 2018 | https://www.encodeproject.org |
| Common TAD boundaries of five human cell-lines | Akdemir et al. 2020 | N/A |
| TRRUST2.0 | Han et al. 2018 | http://www.grnpedia.org/trrust |
| ENCODE Transcription Factor Targets | ENCODE Project Consortium, 2011 | https://maayanlab.cloud/Harmonizome/dataset/ENCODE+Transcription+Factor+Targets |
| Software and Algorithms |  |  |
| FusionAI software | This study | https://ccsm.uth.edu/FusionGDB/FusionAI/ |
| FusionAI model training | This study | https://ccsm.uth.edu/FusionGDB/FusionAI/ |
| STAR-fusion | Haas et al. 2017 | https://github.com/STAR-Fusion/STAR-Fusion |
| Arriba | Uhrig et al. 2021 | https://arriba.readthedocs.io/ |

## CONTACT FOR REAGENT AND RESOURCE SHARING

Further information and requests for materials should be directed to and will be fulfilled by the co-corresponding authors, Drs. Xiaobo Zhou and Pora Kim.

## EXPERIMENTAL MODEL AND SUBJECT DETAILS

None.

## METHOD DETAILS

### FusionAI architecture and training using deep learning (including creation of fusion-negative data)

To train the FusionAI DNN, we used the fusion gene breakpoint information of 48K of fusion genes from FusionGDB. Since most of the fusion genes are predicted from the mRNA level sequencing data and the real genomic breakpoints would be located in intronic regions, we input the exon junctional breakpoints of the known fusion gene breakpoints to train the FusionAI model. Out of ~ 48K known fusion events, about 26K of events had the breakpoints located at the exon junction-junction position (j-j BP combination). To make fusion negative breakpoints data, we excluded 17,110 genes, which is involved in 48K known human fusion genes, among 43K GENCODE genes. From the rest of those genes (27,116), which are not known as involved in any fusion genes, we randomly chose two genes. Then, we filtered out false-positive cases from the unnecessary multiply mapped cases and breakpoints belong to the repeat region, paralogs, or pseudogenes using RepeatMasker, Duplicated Genes Database, and HUGO database’s pseudogenes. This is the typical pre-process by the fusion prediction tools to filter out false positives. We also excluded the gene pairs with neighboring gene relationships to exclude the potential read-through cases. In the case of the intra-chromosomal fusion genes, we set the minimum distance as 100Kbp between randomly chosen two breakpoints across gene bodies. A 20Kbp long DNA sequence was constructed by conjugating +/- 5Kbp sequence from each BP of two partner genes. Through this procedure, we created ~ 26K fusion-negative breakpoints data. Based on the primary sequences, we trained a multiple-layer deep neural network to predict the likelihood of being a fusion gene for a designated gene pair and to identify sequence patterns around the human fusion gene breakpoint area. We designed the input sequence to be +/- 5kb flanking the fusion gene breakpoints of each fusion gene partner gene and output the probability of being fusion and non-fusion. The input of the model is a sequence of 20 kb one-hot encoded nucleotides. The output is two probabilities corresponding to fusion-positive and -negative breakpoint that sum to one. Our deep neural network consists of two convolutional layers with filter size (20,4) and (200,1), one max pooling, one flatten, and two dense layers preceding the output layer. The model involves 1,672,290 parameters including both weight matrix and bias at related layers (Figure S1). 36.4K BPs from a combined total of 52K BPs (26K j-j combination BPs and 26K non-FGBPs) were used as training and validation sets (80% for training and 20% for validation), and the rest 15.6K BPs was used for an independent test. The performance (accuracy and loss) during the training process is illustrated in Figure S2. We then tested the trained model on both the 26K original training samples and the 15.6K test samples. The accuracies for training and test data sets were 97.4% (AUCROC=0.9962) and 90.8% (AUCROC=0.9706) with 0.12 and 0.42 error rate, respectively. This performance is much better than the traditional machine learning method, SVM, that yielded the accuracy of 79% and 72% for training and test data, respectively (Figure 1D).

### Creating simulation RNA-seq data of training and test data to run STAR-fusion and Arriba

We made the simulation RNA-seq data of the split reads at the exon junction breakpoints with different read length (50, 75, and 100 bp) and a different number of split reads (1, 3, 5 split reads, and 10 random around breakpoints) based on the fusion-positive and -negative breakpoints in training and test data sets. Using random module of python, we chose random number once, three, and five times based on the seed length of 25bp as the transcript’s broken position with 0 to 5 varied distance between the 5’-genes’ exon sequence to the breakpoint and the 3’-genes’ exon sequence from the breakpoint to make the split reads at the exon junction site with different read length. We also made 10 random split read sequences with a 10bp distance gap among the read alignments.

### Model evaluation on ChiTaRS-3.1

We downloaded the fusion gene information from ChiTaRS-3.1 (Gorohovski et al., 2017). Among these, we only used the validated fusion genes by the Sanger sequencing approaches, which are stored from the Entrez transcript database by the National Center for Biotechnology Information (NCBI). Among these, 862 fusion genes had the breakpoints at the exon junction-junction positions. For these cases, we made a 20kb long DNA sequence as the input of FusionAI and ran it. Furthermore, to be run these fusion gene data with STAR-fusion and Arriba with default option using GENCODE v19 human genome, we made the simulation RNA-seq data of the split reads at the exon junction breakpoints with different read lengths and the different numbers of split reads.

### Model evaluation on 2,200 fusion genes from 520 cancer cell-lines

We downloaded the fusion gene information of the 2,269 validated in-frame fusion genes from 529 cancer cell-lines by Klijn, C. et al. (Klijn et al., 2015). Out of these, 2,162 fusion genes had the breakpoints at the exon junction-junction. For these cases, we made a 20kb long DNA sequence as the input of FusionAI and ran it. To test STAR-fusion and Arriba, we also made the simulation RNA-seq data with the same way we did for evaluation on ChiTaRS-3.1 data as the input data.

### Comparison with existing fusion gene prediction tools for three cell-lines

K562 is a myelogenous leukemia cell-line with the most famous fusion gene, *BCR-ABL1*. MCF7 is the most studied breast cancer cell-line with multiple identified fusion genes. H2228 is the non-small cell lung cancer cell-line with the *EML4-ALK* fusion gene. We ran STAR-fusion and Arriba for these cell-lines’ RNA-seq data which were downloaded from the Sequence Read Archive (SRA) of NCBI(Leinonen et al., 2011) with SRA accession of SRR521460, SRR064286, and DRR016705.1 for K562, MCF7, and H2228, respectively. To run FusionAI, we made 20kb long DNA sequences based on the breakpoints that were predicted by STAR-fusion and Arriba. For validation, we also made FusionAI input data for the experimentally validated fusion genes among these three cells from the work by Klijn, C. et al.

### Identification of DNA sequence motif

First, we assembled the sequence of the top 10% FI scored regions into the merged sequence contigs as the fasta format when there was a continuous selection of the genomic location as the high FI scored regions. For these unique sequences, we used MEME Suite (Bailey et al., 2009) with the ‘any number of repetitions’ option to identify the enriched DNA sequence motifs around the broken regions by fusion genes per our interests such as all fusion-positives, intra-chromosomal fusion-positives, kinase fusion genes, and transcription factor fusion genes.

### Making FusionAI input data

FusionAI runs based on the primary sequence composed of two genes involved in a fusion gene with +/- 5kb sequence from each breakpoint. For the user input data of two breakpoints of potential fusion genes, the FusionAI package makes the input sequence data using the nibFrag based on the human genome sequence of the hg19 version, which can be downloaded from the UCSC Genome Browser. The tab-delimited data format with fusion gene pairs, chromosome, breakpoint, strand, and input sequence is read by FusionAI model and FusionAI gives the prediction score. In the FusionAI package, the user can make the input data using Run_FusionAI.py based on the interested breakpoint information.

### Human genomic features information

We integrated a total of 44 different types of human genomic feature loci information across five big categories including virus integration sites, repeats, structural variants, chromatin states, and gene expression regulation. First, we downloaded the virus integration site information from the VISDB(Tang et al., 2020) and we lifted it over to the hg19 version using the liftover tool from the UCSC Genome Browser since FusionAI’s training was done based on the sequence of hg19 version (Navarro Gonzalez et al., 2021). We integrated 13 types of repeats (Alu repeats, A-Phased repeats, Directed repeats, DNA transposons, “G-Quadruplex, forming repeats”, Inverted repeats, L1 repeats, L2 repeats, “Low_complexity, A/T rich regions”, Microsatellites, MIR repeats, Mirror repeats, and Z-DNA motifs) from RepeatMasker (Bao et al., 2015) and MicroSatellite DataBase (MSDB) (Avvaru et al., 2020). For the diverse types of structural variants including the copy number variants, we downloaded the arranged breakpoint information of the structural variants from dbVar (Lappalainen et al., 2013). The chromatin states category include the loci of 15 different types of chromatin states such as 1_TssA, 2_TssAFlnk, 3_TxFlnk, 4_Tx, 5_TxWk, 6_EnhG, 7_Enh, 8_ZNF_Rpts, 9_Het, 10_TssBiv, 11_BivFlnk, 12_EnhBiv, 13_ReprPC, 14_ReprPCWk, and 15_Quies, from the previous study on the chromatin state calls using a 15-state model for 12 cell lines, were obtained from the Roadmap Epigenomics Mapping Consortium (Ernst and Kellis, 2017; Roadmap Epigenomics et al., 2015). The gene expression regulatory category includes five types of features as CPGisland, Methylation, Promoters, ReplicationTiming, and TAD boundaries. The information of the first three feature categories was downloaded from the FANTOM5 collection(Lizio et al., 2019). We downloaded the replication timing-specific peak regions from the ENCODE portal site by selecting the assay type of the replication timing (Davis et al., 2018). We used 2,477 loci of common TAD boundaries from a previous study that made high-resolution chromosome conformation (Hi-C) datasets from five human cell lines based on the (Akdemir et al., 2020). The detailed statistics for individual feature categories with their significance results by Fisher’s exact test are in Table S5.

### Gene ontology enrichment analysis

To identify the enriched biological processes in the overlapped genes between the top 10% FI scored regions and individual human genomic features of 44 categories, we used ToppFun of the ToppGene Suite(Chen et al., 2009). We limited the results by the Benjamini and Hochberg false discovery rate < 0.05 and the number of genes in the gene group < 500. We showed the top enriched GO biological process of each feature category in Figure 4B.

## DATA AND SOFTWARE AVAILABILITY

The training and test data and FusionAI model codes, and simulation RNA-seq data are available on https://ccsm.uth.edu/FusionGDB/FusionAI/.

---

**Figure S1. Detailed description of the FusionAI architectures.** FusionAI architecture use flanking nucleotide sequence of length 5,000 on each exon junction breakpoint of interest as input and output the probability of the position being a fusion gene breakpoint or not.

**Figure S2. Training and validation performance for the first 12 epochs.** Shown are the categorical cross entropy loss and overall accuracy, which is the percentage of gene pairs that were correctly classified for both training and test data sets.

